# The fission yeast SUMO-targeted Ubiquitin Ligase Slx8 functionally associates with clustered centromeres and the silent mating type region at the nuclear periphery

**DOI:** 10.1101/2024.09.10.612319

**Authors:** Shrena Chakraborty, Joanna Strachan, Kamila Schirmeisen, Laetitia Besse, Eve Mercier, Karine Fréon, Haidao Zhang, Ning Zhao, Elizabeth H Bayne, Sarah AE Lambert

## Abstract

The SUMO-targeted Ubiquitin ligase (STUbL) family is involved in multiple cellular processes via a wide range of mechanisms to maintain genome stability. One of the evolutionarily conserved functions of STUbL is to promote changes in the nuclear positioning of DNA lesions, targeting them to the nuclear periphery. In *Schizossacharomyces pombe*, the STUbL Slx8 is a regulator of SUMOylated proteins and promotes replication stress tolerance by counteracting the toxicity of SUMO conjugates. In order to study the dynamic dialectic between Ubiquitinylation and SUMOylation in the nuclear space of the *S. pombe* genome, we analyzed Slx8 localization. Unexpectedly, we did not detect replication stress-induced Slx8 foci. However, we discovered that Slx8 forms a single nuclear focus, enriched at the nuclear periphery, which marks both clustered centromeres at the spindle pole body and the silent mating type region. The formation of this single Slx8 focus requires the E3 SUMO ligase Pli1, poly-SUMOylation and the histone methyl transferase Clr4 that is responsible for the heterochromatin histone mark H3-K9 methylation. Finally, we established that Slx8 promotes centromere clustering and gene silencing at heterochromatin domains. Altogether, our data highlight evolutionarily conserved and functional relationships between STUbL and heterochromatin domains to promote gene silencing and nuclear organization.

**Highlights:**

- The *S. pombe* STUbL Slx8 forms a single nuclear focus enriched at the nuclear periphery in a SUMO-chain-dependent manner.
- Slx8 foci mark clustered centromeres and the silenced mating type region but not telomeres.
- H3-K9 methylation by Crl4 promotes the single nuclear Slx8 focus
- Slx8 promotes centromere clustering and gene silencing.

## Introduction

The nuclear architecture and the 3D genome organization have emerged as important regulation layers of genome maintenance, contributing to numerous DNA-associated transactions such as chromosome segregation, transcription and DNA repair (Misteli and Soutoglou, 2009). Chromatin displays functional compartmentalization: while gene-rich, transcriptionally active chromatin tends to localize to the interior of the nucleus, gene-poor, transcriptionally repressed heterochromatin is typically enriched at the nuclear periphery (NP), which is believed to provide a microenvironment favoring association of factors required for silencing (reviewed in (Towbin et al., 2009)). In many organisms, centromeres also cluster together at the NP, and this spatial organisation has been shown to be important for promoting loading of centromeric proteins (Wu et al., 2022), silencing of repetitive elements (Padeken et al., 2013), and the prevention of micronuclei formation (Jagannathan et al., 2018). The stability of the genome is particularly vulnerable during the process of DNA replication since a broad spectrum of obstacles can jeopardize the progression of the replication machinery, resulting in fork stalling, collapse or breakage (Zeman and Cimprich, 2014). In several organisms, from yeast to flies and mammalian cells, DNA lesions, including double strand break (DSB) and replication stress site, shift away from their initial nuclear compartment to associate with the NP. Such mobility of DNA lesions allows a spatial regulation of DNA repair processes to ensure optimal error-free repair outcome (reviewed in (Lamm et al., 2021; Whalen and Freudenreich, 2020).

The NP is composed of a double membrane nuclear envelop (NE) and multiple nuclear pore complexes (NPCs) embedded in the NE. In yeast, the spindle pole body (SPB), the functional macromolecular structure equivalent to centrosome, is also embedded in the NE. Components of both the NE and the NPC have been reported as factors allowing anchorage of DNA lesions to the NP (reviewed in (Whalen and Freudenreich, 2020). Although the mechanisms of relocation and anchorage differ depending on the type of DNA lesion and the cell cycle stage, an emerging common feature is the requirement for SUMOylation which homeostasis is critical to maintain genome integrity (Schirmeisen et al., 2021). SUMO (small ubiquitin-like modifier) is a post-translational modification present in all eukaryotic systems. SUMO is covalently attached to a target thanks to the coordinated activity of E2 and E3 SUMO ligases (reviewed in (Chang et al., 2021)). Target proteins can be either mono-SUMOylated on a single lysine residue, or harbor multiple single SUMO modifications on several lysine residues, a type of poly-SUMOylation. Moreover, additional SUMO molecule can be covalently attached to the internal lysine of SUMO to form SUMO chains, another type of poly-SUMOylation. SUMOylation affects the activity, the localization and stability of modified targets, with SUMO chains often favoring protein degradation.

A key determinant of the fate of SUMOylated proteins is the SUMO-targeted E3 Ubiquitin ligase (STUbL) family that recognizes SUMOylated proteins and attaches ubiquitin to them. STUbLs are involved in diverse molecular processes, including DNA repair and replication, both during unchallenged conditions and in response to genotoxic stresses (reviewed in (Chang et al., 2021)). STUbls are characterized by a RING-type E3 ubiquitin ligase domain and one or several SUMO-interacting motifs (SIMs) to recognize SUMOylated substrates. Modification by STUbLs can target substrates for proteosomal degradation or mediate non-proteolytic functions. STUbLs act in specific environments, such as the NE, centromere, kinetochore or PML nuclear bodies in human cells. STUbLs have also been implicated in localizing DSBs and replication stress sites to the NP to promote DNA repair and fork restart (reviewed in (Lamm et al., 2021; Whalen and Freudenreich, 2020)). A seminal study in *Saccharomyces cerevisiae* (Sc) first showed that difficult-to repair DSBs and collapsed forks anchor to the NPC in a process requiring the ScSlx5-Slx8 STUbL that physically associates with the Nup84 complex, a component of the NPC (Nagai et al., 2008). Further studies established that the SUMOylation status of proteins bound to DSBs influences the target destination. For example, mono-SUMOylation allows S-phase DSBs to relocate to Mps3, a NE component, whereas poly-SUMOylation allows DSBs in G1 to associate with the NPC in STUbL-dependent manner, suggesting a specificity of STUbL for poly-SUMO chains (Horigome et al., 2014; Horigome et al., 2016).

The target destination of replication stress sites described so far is the NPC. This includes forks stalled within telomeres sequences, at tri-nucleotides repeats, at a replication fork barrier (RFB) mediated by DNA-bound protein and forks stalled by global replication stress in human cells (Aguilera et al., 2020; Kramarz et al., 2020; Nagai et al., 2008; Pinzaru et al., 2020; Rivard et al., 2024; Su et al., 2015). In *S. cerevisiae*, forks stalled at expanded CAG repeats, anchor to the NPC in a process that requires the SIMs of Slx5 and mono-SUMOylation, since preventing poly-SUMOylation does not affect relocation to the NP (Su et al., 2015). The SUMOylation of the repair factors RPA, Rad52 and Rad59 is sufficient to trigger Slx5-dependent relocation to the NP, suggesting that Slx5 may recognize several SUMO particles covalently attached to distinct targets (Whalen et al., 2020). Targeting forks stalled at CAG repeats to the NPC allows the loading of the recombinase Rad51 and prevents the chromosomal fragility of CAG repeats.

In *Schizosaccharomyces pombe*, we have revealed a SUMO-based mechanism that allows the spatial regulation of the recombination-dependent replication (RDR) process, a mechanism that ensures the restart of arrested forks by homologous recombination (Kramarz et al., 2020). Forks arrested by the *RTS1*-RFB relocate to the NP to associate with the NPC in a process requiring SUMO chain formation and the SpSTUbL. In *S. pombe*, Rfp1 and Rfp2 are functional homologs of ScSlx5 but lack E3 activity. They recruit Slx8 through a RING-RING domain interaction to form a functional E3 Ubiquitin ligase (Prudden et al., 2007; Prudden et al., 2011). The absence of a functional spSlx8 STUbL results in the accumulation of high molecular weight (HMW) SUMO conjugates and sensitivity to genotoxic drugs that can be alleviated by the inactivation of the E3 SUMO ligase Pli1 and by preventing SUMO chain formation, suggesting that SpSTUbL has a specificity in targeting poly-SUMOylated substrates (Kosoy et al., 2007; Nie et al., 2017; Prudden et al., 2007; Steinacher et al., 2013). We further established that the relocation of the RFB to the NP promotes RDR via two activities that are enriched in the NPC environment, namely the SUMO protease Ulp1 and the proteasome (Schirmeisen et al., 2024).

One of the unresolved questions in the field is to understand the dynamic crosstalk between SUMOylation and Ubiquitination during the process of relocation of stressed forks and how such crosstalk is spatially segregated in the nuclear space. For example, both SUMOylation and STUbL activity are expected to occur at the site of replication stress before relocation to the NP. Indeed, the drosophila STUbL Dgrn (for degringolade) is recruited at heterochromatic DSBs prior to relocation and after the action of E3 SUMO ligases (Ryu et al., 2015; Ryu et al., 2016). To investigate the temporal and spatial dynamics of SpSlx8 by live cell imaging in response to global replication stress, we generated a functional fusion protein Slx8-GFP, in a similar approach to the one employed to characterize damage-induced ScSlx5 foci (Cook et al., 2009) and SpUfd1 (for ubiquitin-fusion degradation protein) that physically interacts with STUbL (Køhler et al., 2013). We observed that Slx8-GFP did not form replication-stress induced foci but a single discrete focus enriched at the NP in unstressed condition. Both SUMO chains and the E3 SUMO ligase Pli1 are necessary to sustain Slx8-GFP focus formation. Further cellular analysis established that Slx8-GFP focus marks heterochromatin domains positioned at the NP and in the SPB environment, including centromeres and the mating type (*mat*) region. Both heterochromatin and anchoring of centromeres to SPB promotes Slx8-GFP focus. Finally, we provide functional evidence that Slx8 is actively involved in gene silencing and in the clustering of centromeres. Our results highlight functional and physical crosstalk between STUbL and heterochromatin to orchestrate the nuclear organization of specific domains.

## Results

### Slx8-GFP forms a single nuclear focus in a SUMO chain dependent manner

To investigate the spatial dynamics of SUMO conjugates prone to STUbL-dependent processing, Slx8 was C-terminally tagged with GFP, and Slx8-GFP functionality was established based on resistance to genotoxic stress (Figure 1A-B). To further confirm that the GFP tag did not interfere with Slx8 function, we analyzed global SUMO conjugates by immuno-blot. We observed an accumulation of high molecular weight (HMW) SUMO conjugates in the strain bearing the temperature-sensitive *slx8-29* allele when grown at the restrictive temperature (35°C), but not at the permissive temperature (25°C), indicating defective processing of SUMO conjugates in the absence of a functional Slx8 pathway, as expected (Figure 1C) (Nie et al., 2017). None of these HMW SUMO conjugates were detected in WT or Slx8-GFP expressing strains in untreated conditions, whereas they accumulated similarly in both strains upon cells exposure to methyl methane sulfonate (MMS), an alkylating agent known to induce global SUMOylation (Figure 1C) (Nie et al., 2017). These results confirm that the Slx8-GFP fusion protein is functional. Then, we performed live cell imaging and observed that Slx8-GFP formed a single bright focus in most septated cells, which correspond to the bulk of S-phase, and mono-nucleated cells, which mainly correspond to G2 cells (Figure 1 D-E). To address the link between this single Slx8-GFP focus and SUMO metabolism, we investigated the role of the two E3 SUMO ligases known in *S. pombe*: the SUMO chain-modified Pli1 and Nse2 (Andrews et al., 2005; Steinacher et al., 2013). We made use of point mutations in the RING domain of each protein to abolish the E3 SUMO ligase activity. Global SUMOylation was considerably reduced in cells expressing the mutated form Pli1-RING^mut^, compared to WT, and no MMS-induced SUMO conjugates were detected (Figure 2A), consistent with Pli1 being responsible for most of global SUMOylation. In contrast, the global level of SUMO-conjugates was unaffected in cells expressing the mutated form Nse2-RING^mut^, despite this mutation rendering cells sensitive to genotoxic agents (Figure 2A and Figure S1A). Of note, the combination of Slx8-GFP with either Pli1-RING^mut^ or Nse2-RING^mut^ did not aggravate the cell sensitivity to genotoxic agents, further confirming the functionality of Slx8-GFP (Figure S1A). Interestingly, the Slx8-GFP focus was less frequently observed in S and G2-phase of *pli1-RING^mut^* cells, whereas no differences were detected in *nse2-RING^mut^* cells, compared to WT (Figure 2B-C). Of note, the expression level of Slx8-GFP in *pli1-RING^mut^* and *nse2-RING^mut^* was similar to WT, excluding that the lack of Slx8-GFP focus resulted from an expression defect (Figure S1B-C). We were unable to address the potential overlapping role of Nse2 and Pli1 in promoting Slx8-GFP focus formation, since spores harboring both *pli1-RING^mut^* and *nse2-RING^mut^* alleles were unviable. We concluded that the SUMO E3 ligase Pli1, that is responsible for global SUMOylation, sustains the formation of the single nuclear Slx8-GFP focus.

**Figure 1:**
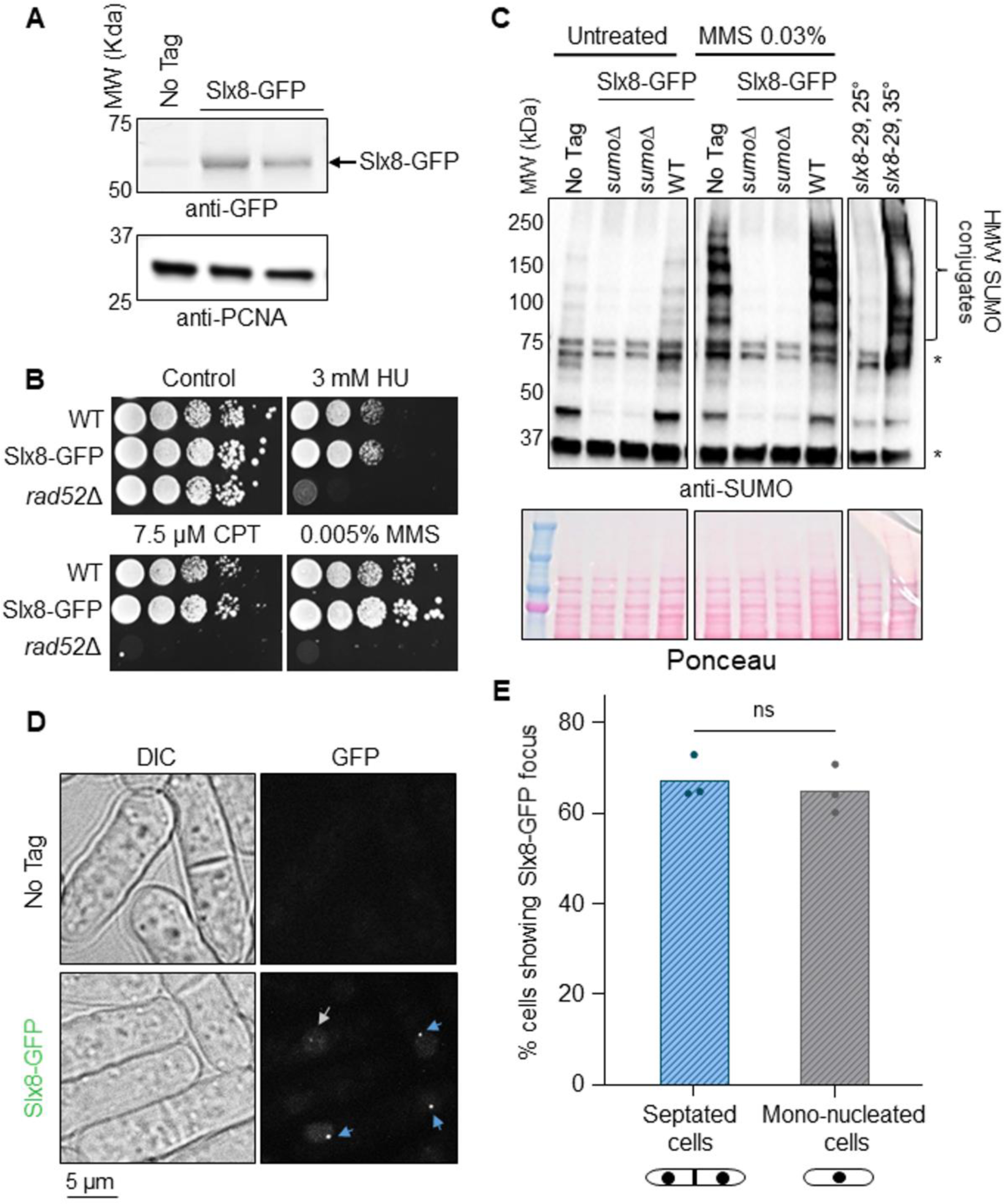
Slx8-GFP forms a single focus in unstressed conditions. **A.** Expression of the endogenously GFP-tagged Slx8 fusion protein. An untagged WT strain (No Tag) was included as control for antibody specificity. PCNA was used as a loading control. Slx8-GFP has a molecular weight (MW) of 58 KDa. **B.** Sensitivity of indicated strains to indicated genotoxic drugs. Ten-fold serial dilution of exponential cultures were dropped on appropriate plates. HU: hydroxyurea; CPT: camptothecin and MMS: methyl methane sulfonate. **C.** Expression of SUMO conjugates in indicated strains and conditions. A strain deleted for *pmt3* gene that encodes the SUMO particle (*sumo*Δ) was added as control for antibody specificity. * indicates unspecific signal. A strain bearing the temperature-sensitive allele *slx8-29* was grown at permissive (25°C) and restrictive (32°C) temperature. **D.** Example of bright-field (left panel, DIC) and GFP fluorescence (right panel) images of cells expressing the endogenous Slx8-GFP fusion protein in indicated strains. Blue and white arrows indicate Slx8-GFP foci in septated and mono-nucleated cells, respectively. Scale bar is 5 µm. **E.** Histogram plots showing the percentage of septated and mono-nucleated cells with nuclear Slx8-GFP foci. *p* value was calculated by two-tailed t test (ns: non-significant). Dots represent values obtained from three independent biological experiments. At least 200 nuclei were analyzed for each strain and cell type.

**Figure 2:**
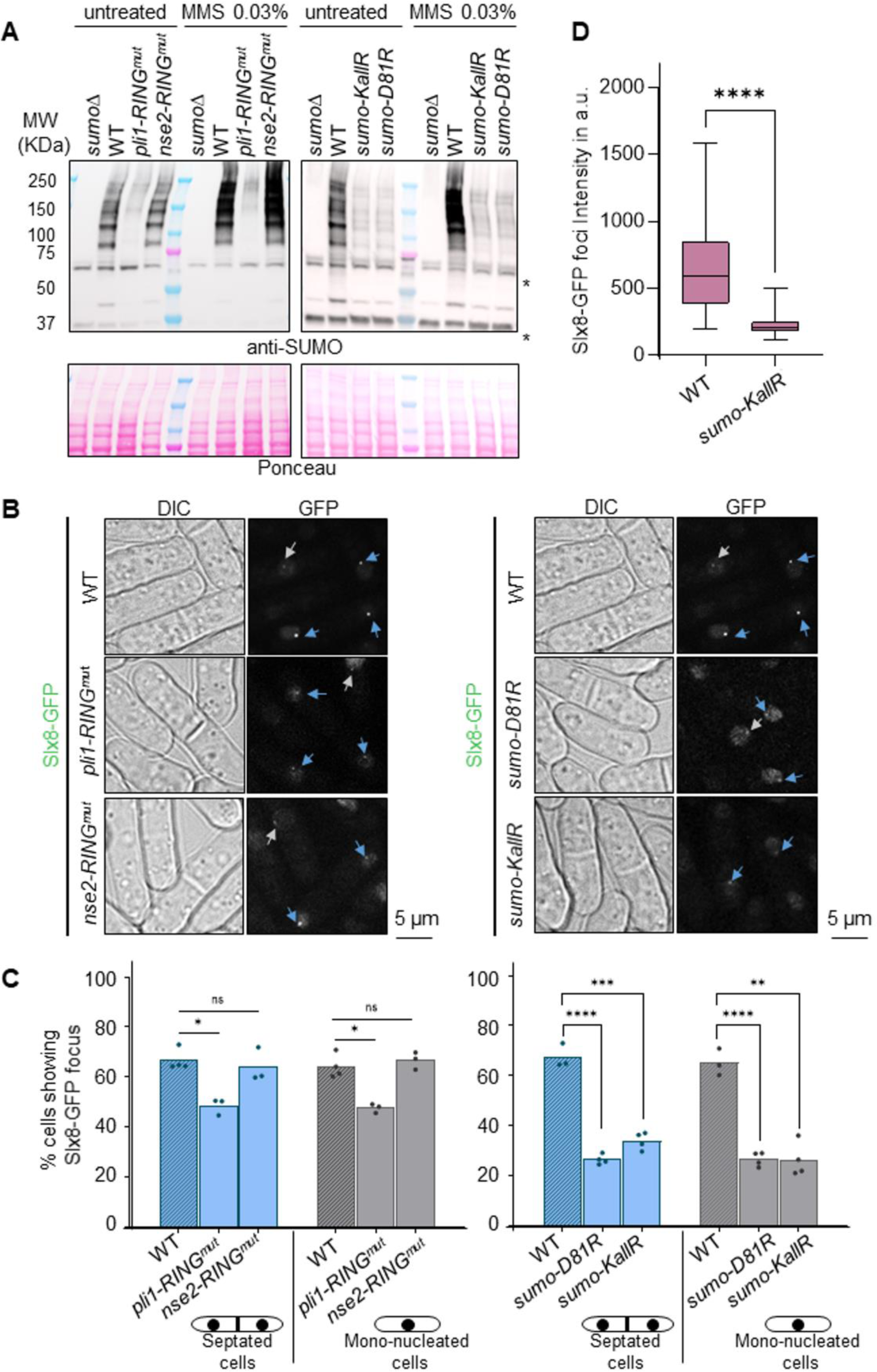
SUMOylation promotes the formation of Slx8-GFP foci. **A.** Expression of SUMO conjugates in indicated strains (expressing Slx8-GFP) and conditions. A strain deleted for *pmt3* gene that encodes the SUMO particle (*sumo*Δ) was added as control for antibody specificity. * indicates unspecific signal. **B.** Example of bright-field (DIC) and GFP fluorescence images in indicated strains expressing Slx8-GFP. Blue and white arrows indicate Slx8-GFP foci in septated and mono-nucleated cells, respectively. Scale bar is 5 µm. **C.** Histogram plots showing the percentage of septated and mono-nucleated cells with nuclear Slx8-GFP foci in indicated strains. *p* value was calculated by two-tailed t test (**** *p* ≤0.0001; *** *p*≤0.001; ** *p*≤0.01; * *p*≤0.05; ns: non-significant). Dots represent values obtained from independent biological experiments. At least 200 nuclei were analyzed for each strain and cell type. **D.** Box-and-whisker plots of Slx8-GFP intensity (mean fluorescence intensity) in indicated strains. Boxes represent the 25/75 percentile, black lines indicate the median, the whiskers indicate the 5/95 percentile. *p* value was calculated by Mann-Whitney U test (**** *p* ≤0.0001). Values were obtained from at least two independent biological experiments. At least 60 nuclei were analyzed for each strain.

Next, we asked which type of SUMOylation contributes to the formation of the Slx8-GFP focus. We could not employ the strain harboring the deletion of the SUMO particle (*pmt3Δ*, here after SUMOΔ), since this strain is extremely sick, showing frequent nuclear deformation. Instead, we employed a strain expressing SUMO-KallR, in which all internal lysine are mutated to arginine to prevent SUMO chain formation (Kramarz et al., 2020) and a strain expressing SUMO-D81R that allows mono and di-SUMOylation to occur but impairs the chain-propagating role of Pli1 (Prudden et al., 2011). As expected, global SUMOylation was massively reduced in strains expressing SUMO-KallR and SUMO-D81R, even upon MMS treatment, compared to WT (Figure 2A). Consistently, the frequency of cells showing a single nuclear Slx8-GFP focus was reduced by almost two-thirds in SUMO-KallR and SUMO-D81R cells, compared to WT (Figure 2B-C), indicating that SUMO-chains are critical determinants of Slx8-GFP focus formation. Of note, Slx8-GFP expression level was only slightly reduced (by ∼ 20%) in SUMO-D81R, an insufficient reduction to explain the lack of two-thirds of the foci (Figure S2). In addition to being less frequently formed, Slx8-GFP foci were three to four times less intense in SUMO-KallR cells, compared to WT (Figure 2D). We concluded that the formation of the single nuclear Slx8-GFP focus requires SUMO chain formation and the SUMO-chain modified E3 ligase Pli1, suggesting that it marks SUMO conjugates at specific nuclear regions.

### Slx8-GFP does not form supernumerary foci in response to replication stress

Having establish that Slx8-GFP marks specific nuclear regions in a SUMO-dependent manner, we investigated if slx8-GFP forms DNA damage-induced foci, as reported for ScSlx5 (Cook et al., 2009). Treatment with MMS, but not with hydroxyurea (HU, an inhibitor of the ribonucleotide reductase leading to a depletion of the dNTP pool and stalled replication fork), or camptothecin (CPT, an inhibitor of the topoisomerase I leading to collapsed replication fork), resulted in a marked accumulation of SUMO conjugates (Figure 3A). Whatever the replication-blocking agent used, no additional DNA damage-induced Slx8 foci could be detected in our microscopy setup on living cells, even in condition of MMS-induced accumulation of SUMO conjugates (Figure 3A-C). Surprisingly, HU treatment resulted in a 50% reduction in cells showing a single Slx8 focus in WT cells. It’s worth noting that, despite the absence of supernumerary Slx8-GFP foci, the intensity of the single Slx8-GFP focus increased significantly upon exposure to genotoxic stresses, particularly after MMS treatment, compared with the untreated condition (Figure 3D). Furthermore, the frequency of cells showing a single Slx8-GFP focus was severely reduced in SUMO-KallR cells after treatment with replication blocking agents (Figure 3B-C) suggesting that SUMO chains become more critical for maintaining the Slx8 GFP focus under replication stress conditions. Although we observed a slight decrease in Slx8-GFP expression in WT and SUMO-KallR cells in response to treatments (Figure S3), the extent of variation seems insufficient to explain the disappearance of Slx8-GFP foci. We concluded that Slx8-GFP cannot serve as a readout of damage-induced SUMO chain formation but that the behavior of the single Slx8-GFP focus is modulated by replication stress in a SUMO-chain dependent manner.

**Figure 3:**
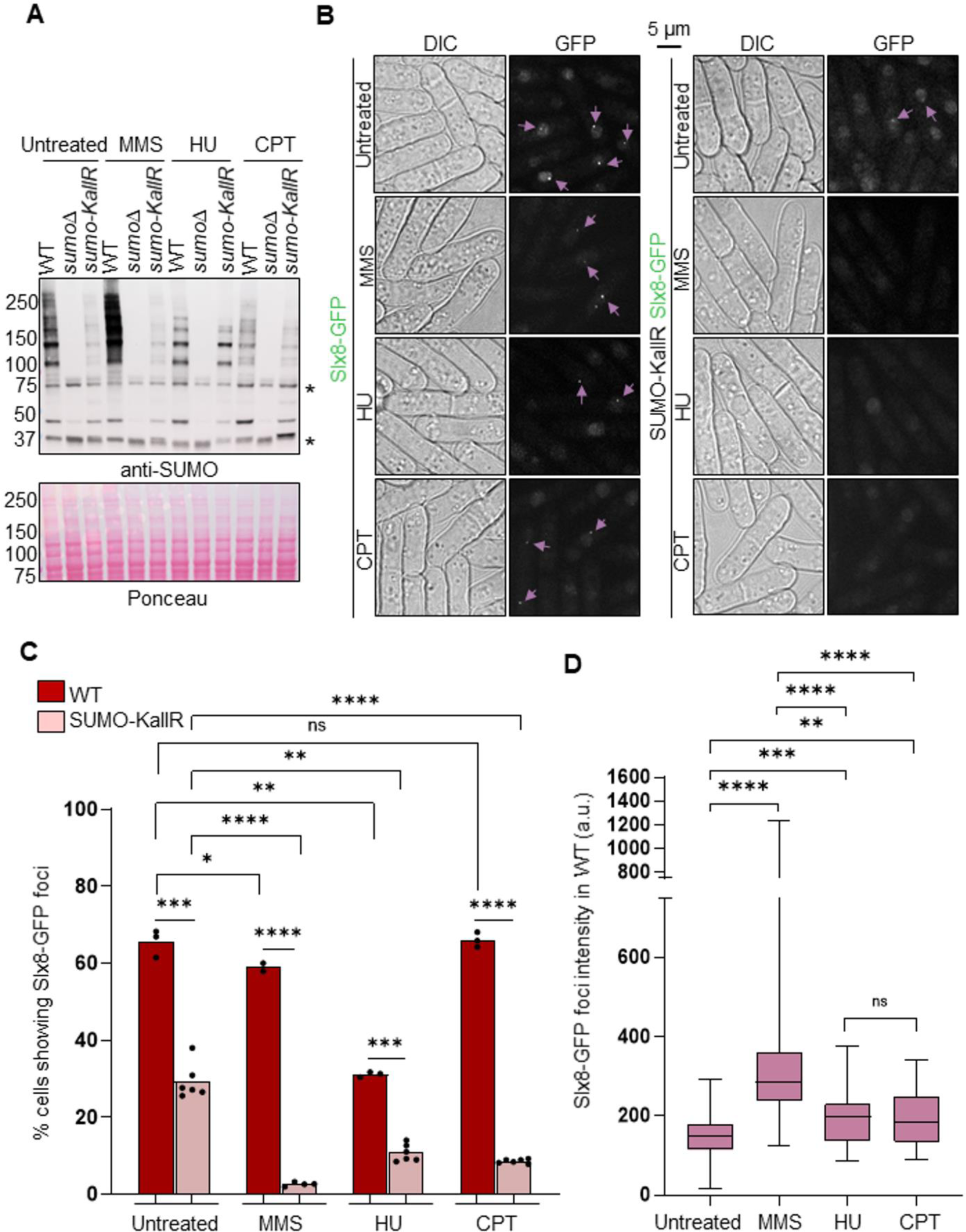
Genotoxic stress does not lead to supernumerary Slx8-GFP foci. **A.** Expression of SUMO conjugates in indicated strains (expressing Slx8-Gfp) and conditions. A strain deleted for *pmt3* gene that encodes the SUMO particle (*sumo*Δ) was added as control for antibody specificity. * indicates unspecific signal. Strains were treated with genotoxic drugs before the extraction of proteins. HU: hydroxyurea (20 mM, 4 hours); CPT: camptothecin (40 µM, 4 hours) and MMS: methyl methane sulfonate (0.03%, 3 hours). **B.** Example of bright-field (DIC) and GFP fluorescence (panel) images in indicated strains and conditions. Genotoxic stresses were generated as in A. Pink arrows indicate cells harboring nuclear Slx8-GFP foci. Scale bar is 5 µm. **C.** Histogram plots showing the percentage of cells with nuclear Slx8-GFP foci in indicated strains and conditions. *p* value was calculated by two-tailed t test (**** *p* ≤0.0001; *** *p*≤0.001; ** *p*≤0.01; * *p*≤0.05; ns: non-significant). Dots represent values obtained from two independent biological experiments. At least 200 nuclei were analyzed for each strain and treatment condition. **D.** Box-and-whisker plots of Slx8-GFP intensity (mean fluorescence intensity) in indicated strains and conditions. Boxes represent the 25/75 percentile, black lines indicate the median, the whiskers indicate the 5/95 percentile. *p* value was calculated by Mann-Whitney U test (**** *p* ≤0.0001; *** *p*≤0.001; ** *p*≤0.01). Values were obtained from two independent biological experiments. At least 60 nuclei were analyzed for each strain and treatment condition.

### The single nuclear Slx8-GFP focus marks centromere and the *mat* region at the nuclear periphery

The analysis of cell images revealed that the single Slx8-GFP focus in untreated condition was often positioned at the periphery of the nucleus. To confirm this, we asked how frequently Slx8-GFP foci co-localize with Cut11-mCherry, a component of the NPC that marks the NP. We found that the nuclear Slx8-GFP focus, where visible, was positioned at the NP in ∼ 65 % of WT S-phase cells (septated cells) and this frequency dropped to ∼ 35 % in WT G2 cells (mono-nucleated cells) (Figure 4A-B). Interestingly, this peripheral nuclear positioning in S-phase dropped to ∼ 35 % in cells expressing SUMO-KallR. We concluded that most Slx8 foci are enriched at the NP and that SUMO chains contribute both to Slx8-GFP focus formation and positioning at the NP during S-phase.

**Figure 4:**
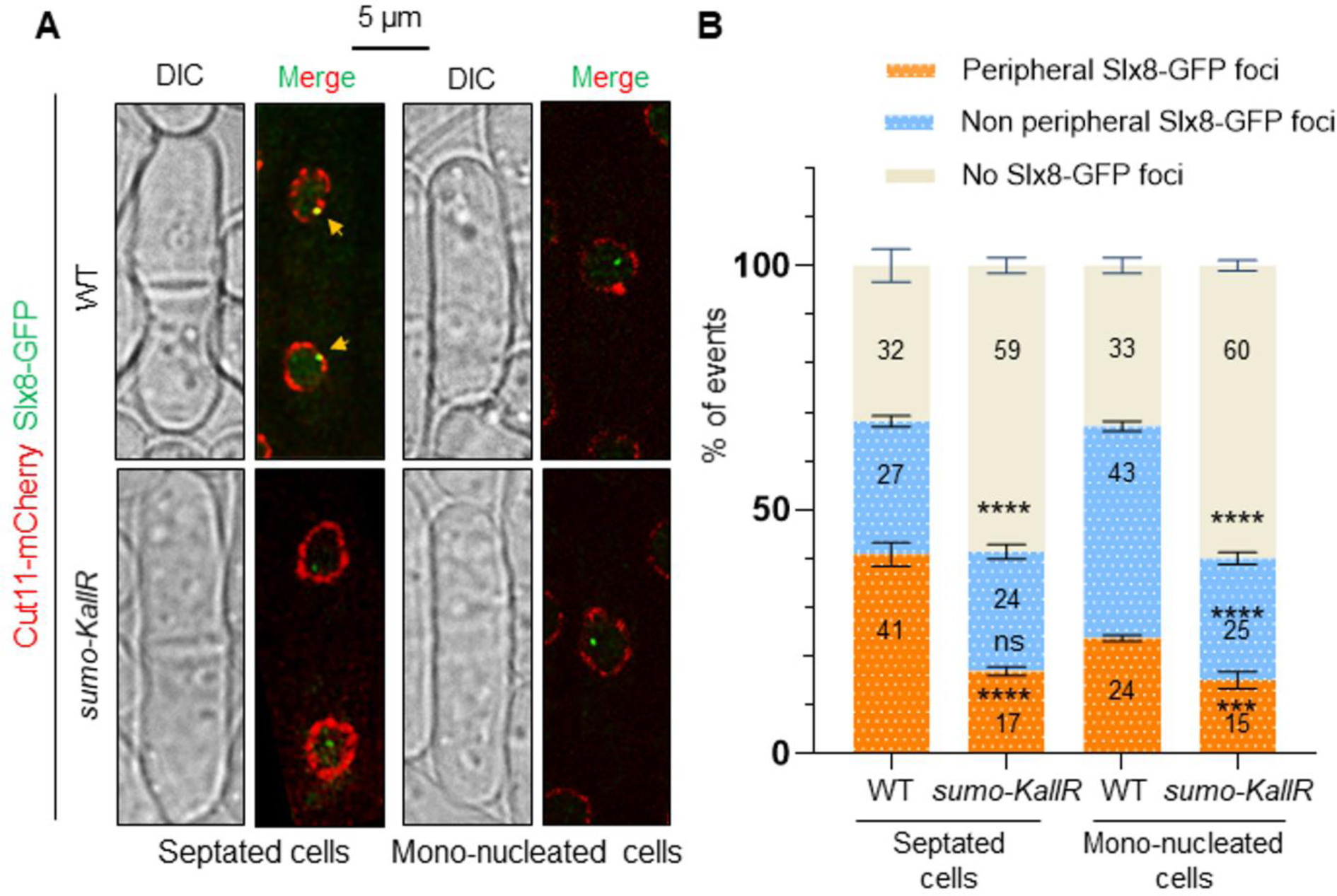
Slx8-GFP focus is enriched at the nuclear periphery. **A.** Representative cell images of cells expressing Cut11-mCherry (red) and Slx8-GFP (green) in septated and mono-nucleated cells of indicated strains. The nuclear periphery is visualized via Cut11-mCherry. Yellow arrows indicate co-localization events. Scale bar is 5 µm. **B.** Stacked bar charts showing the frequency of co-localization between Slx8-GFP and Cut11-mCherry in septated and mono-nucleated cells of indicated strains. Individual bars represent 100% of events and numbers indicate the % of each category (peripheral Slx8-GFP foci co-localizing with Cut11-mCherry in orange, non-peripheral Slx8-GFP foci in blue, absence of Slx8-GFP foci in cream-white). *p* value was calculated by two-tailed t-test (**** *p* ≤0.0001; *** *p*≤0.001; ns: non-significant). Bars indicate mean values ±Standard deviation (SD). Values were obtained from two independent biological experiments. At least 200 nuclei were analyzed for each strain and cell type.

The peripheral nuclear location of the single Slx8-GFP focus suggests that Slx8 associates with specific components and/or chromosomal regions known to be at the NP. During interphase, the *S. pombe* chromosomes are arranged in a Rabl-like configuration in which the three centromeres are clustered adjacent to the SPB embedded in the NE, while telomeres form discrete foci clustered at the NP at the opposing hemisphere of the nucleus (Mizuguchi et al., 2015). In addition, the heterochromatin domain of the sexual mating locus (hereafter *mat* region), that contains the silent *mat2* and *mat3* loci, is also positioned at the NP nearby the SPB. We thus addressed if Slx8-GFP localizes with markers of centromere (Mis6-RFP, a kinetochore component), SPB (Sid4-RFP) and telomere (Taz1-RFP) and the *mat* region (using a strain harboring a *LacO* array integrated nearby the *mat* locus, bound by the fluorescent repressor LacI-mCherry) (Figure 5A). During S-phase (in septated cells), the nuclear Slx8-GFP focus co-localized with Sid4-RFP and Mis6-RFP in ∼ 60 % of cells showing a Slx8-GFP focus, whereas a co-localization event with the *mat* region was observed in ∼ 20 % of the cells (Figure 5B). Such nuclear positioning appeared highly significant compared to random co-localization events. Although less pronounced, the Slx8-GFP focus significantly overlapped with centromere, SPB and *mat* region in G2 cells (mono-nucleated cells). In contrast to the Slx8-GFP focus, all cells exhibited a single Sid4-RFP and Mis6-RFP focus, or a single LacI-mcherry dot marking the *mat* region (Figure 5A). We found that centromere and SPB are positively associated with Slx8 in 40 % of S-phase cells and in 20% of G2 cells, whereas the *mat* region associated with Slx8 in ∼ 15-18 % of S and G2-phase cells (Figure 5C). In contrast, no co-localization above random events were detect between Slx8-GFP and Taz1-marked telomeres foci. We concluded that for the most part, the Slx8-GFP focus positioned at the NP marks clustered centromeres, the SPB and the *mat* region.

**Figure 5:**
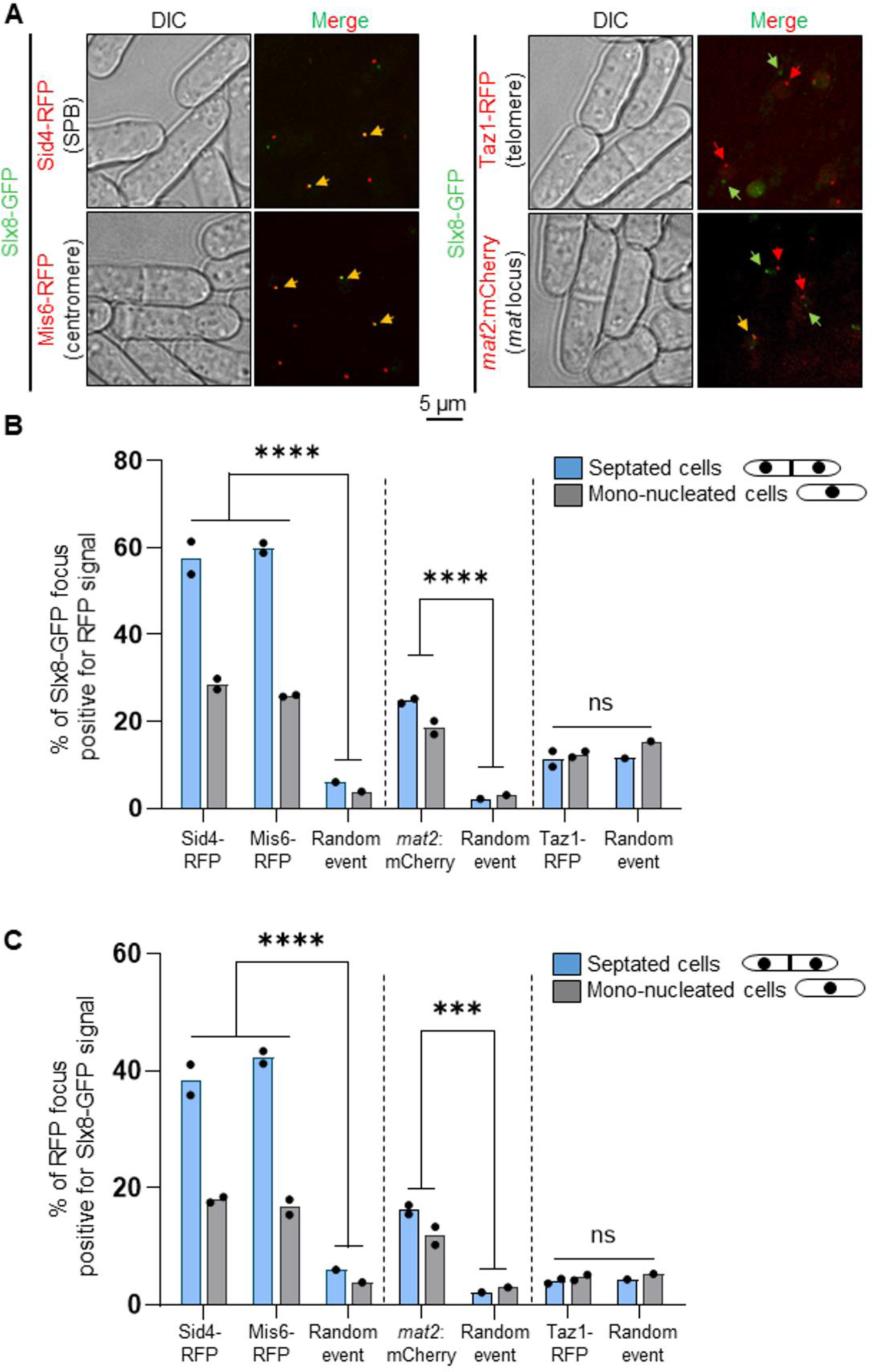
Slx8-GFP focus marks the SPB, centromere and the mating type locus. **A.** Representative cell images of strains expressing Slx8-GFP and either Sid4-RFP (a SPB marker) or Mis6-RFP (a kinetochore marker) or Taz1-RFP (a telomere marker), or harboring the endogenous *mat2* locus tagged with a LacO arrays bound by LacI-Mcherry (*mat2:mCherry*). Red, green and yellow arrows indicate RFP, GFP and co-localization events, respectively. Scale bar is 5 µm. **B & C.** Histogram plots showing the percentage of co-localization events between Slx8-GFP and the above described markers. *p* value was calculated by Brown-Forsythe and Welch ANOVA test (**** *p* ≤0.0001; *** *p*≤0.001; ns: non-significant). Dots represent values obtained from two independent biological experiments. At least 200 nuclei were analyzed for each strain and cell type.

### Heterochromatin and centromeres clustering at SPB sustain Slx8-GFP focus formation

Slx8-GFP marks the SPB environment and associated chromosomal regions such as centromeres and *mat* region, both being enriched for heterochromatin that ensures gene silencing. Therefore, we asked if heterochromatin formation and centromere clustering are required to ensure the formation of a single Slx8-GFP focus. We observed that in the absence of Clr4, the histone methyl-transferase that promotes H3-K9 methylation, a hallmark of heterochromatin and gene silencing (Nakayama et al., 2001; Rea et al., 2000), the frequency of Slx8-GFP focus formation was reduced by two fold (Figure 6A-B). In contrast, no effect was observed in the absence of Dicer (Dcr1), a component of the RNAi machinery promoting the establishment of heterochromatin, but with only a partial role in maintenance. These results indicate that H3K9 methylation, but not RNAi, is required to promote the formation of the nuclear Slx8-GFP focus. We also investigated the role of centromere clustering. Csi1 is a key factor that provides a physical link between kinetochores and SPB associated proteins. The lack of Csi1 leads to a severe defect in centromere clustering (Hou et al., 2012) and resulted in a 2 fold reduction in the frequency of the Slx8-GFP focus (Figure 6A-B). Of note, the expression of Slx8-GFP was not affected in the absence of Csi1, Clr4 or Dcr1, indicating that the decreased in the frequency of Slx8-GFP foci is not caused by variation in expression level (Figure S4). Thus, both heterochromatin formation and centromere clustering contribute to Slx8-GFP focus formation.

**Figure 6:**
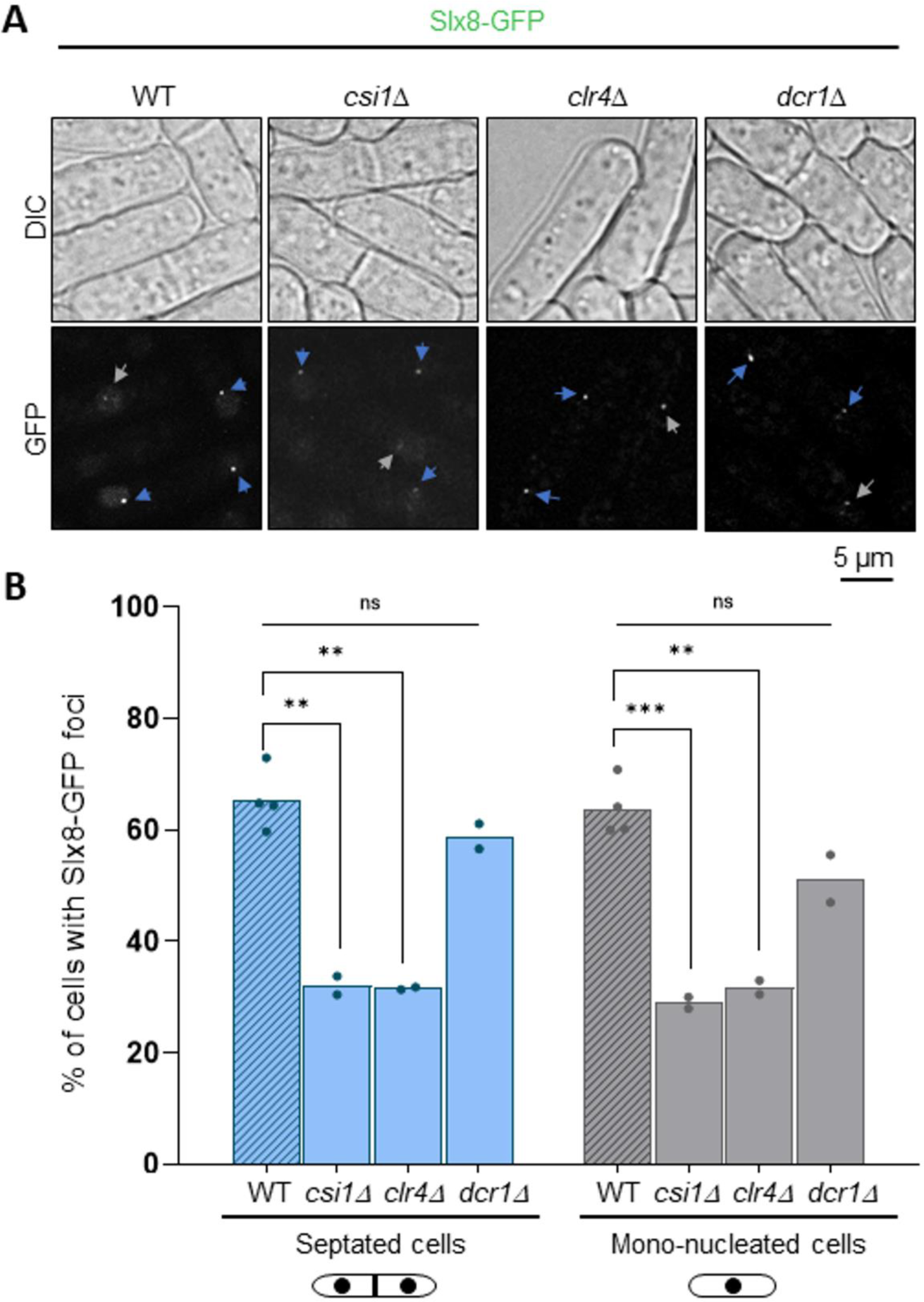
Heterochromatin and centromere clustering promote Slx8-GFp focus formation. **A.** Example of bright-field (top panel, DIC) and GFP fluorescence (bottom panel) images of cells expressing Slx8-GFP in indicated strains. Blue and white arrows indicate Slx8-GFP foci in septated and mono-nucleated cells, respectively. Scale bar is 5 µm. **B.** Histogram plots showing the percentage of septated and mono-nucleated cells with nuclear Slx8-GFP foci. *p* value was calculated by two-tailed t test (*** *p*≤0.001; ** *p*≤0.01; ns: non-significant). Dots represent values obtained from two independent biological experiments. At least 200 nuclei were analyzed for each strain and cell type.

### Slx8 promotes centromere clustering and gene silencing

Finally, we tested whether Slx8 functions to promote heterochromatic silencing and centromere clustering. To assess silencing, we performed RT-qPCR analysis of transcripts from the heterochromatic pericentromere (*cen[dg]*) and silent mating-type (*mat*) regions. Such transcripts accumulate at very low levels in wild-type cells, but much higher levels in absence of factors such as Clr4 required for heterochromatin assembly. Interestingly, we also observed a small but significant increase in accumulation of transcripts from both the pericentromere and the *mat* locus in cells lacking Slx8, consistent with Slx8 functionally contributing to silencing in these regions (Figure 7A). To assess centromere clustering, we performed live-cell imaging on cells expressing GFP–Cnp1 (*S. pombe* CENP-A, the centromere-specific histone variant) to visualise centromeres, together with Sid4–RFP as a marker of the SPB. Whereas wild-type cells consistently display a single GFP–Cnp1 focus, representing three clustered centromeres, adjacent to the SPB, absence of Csi1 results in ∼35% of cells showing more than one GFP–Cnp1 focus, indicative of defective clustering. Strikingly, the lack of Slx8 also resulted in a significant clustering defect, with ∼12% of cells displaying more than one GFP–Cnp1 focus (Figure 7B-C). An epistatic phenotype was seen for *slx8Δ csi1Δ* double mutant cells, which displayed clustering defects comparable to those in the *csi1Δ* single mutant, suggesting that Slx8 may function in the same pathway as Csi1. Deletion of the SUMO ligase Pli1 largely suppressed the clustering defect associated with absence of Slx8, consistent with it arising as a result of excess SUMOylation. We conclude that localization of Slx8 in the vicinity of the SPB both depends on, and contributes to, heterochromatin integrity and centromere clustering.

**Figure 7:**
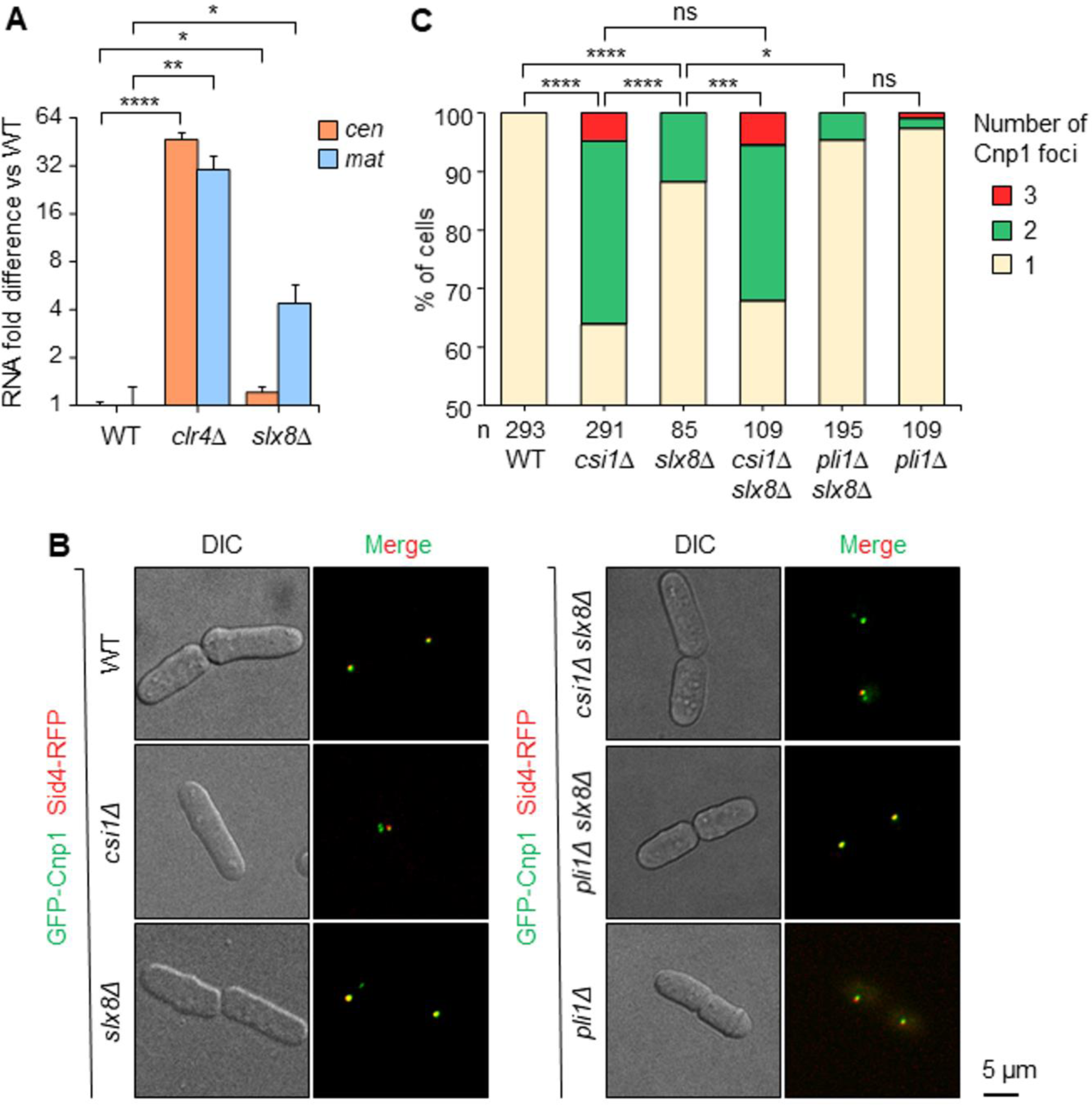
Slx8 promotes heterochromatic silencing and centromere clustering. **A.** RT-qPCR analysis of pericentromere (*cen[dg]*) and mating type locus (*mat*) transcript levels, normalized to control transcript *act1^+^*. Data are plotted as fold difference relative to wild-type, on log_2_ scale. *p* value was calculated by two-tailed t test (**** *p*≤0.0001; ** *p*≤0.01; * *p*≤0.05). Values are mean ±s.d. from three independent experiments. **B.** Representative images of cells expressing GFP–Cnp1 (centromere marker) and Sid4–RFP (SPB marker). Scale bar is 5 µm. **C.** Stacked bar charts showing the percentage of cells displaying one, two or three Cnp1 foci, based on analysis of n nuclei. *p* value was calculated by chi-squared test (**** *p* ≤0.0001; *** *p*≤0.001; * *p*≤0.05; ns: non-significant).

## Discussion

STUbL proteins play diverse roles throughout the cell cycle to protect against genome instability. Here, we revealed that the fission yeast STUbL Slx8 functions mainly in the SPB environment in a SUMO-dependent manner to help ensure centromere clustering and gene silencing at heterochromatic domains. These data are consistent with SUMOylation of centromeres being an important mediator of centromere identity, and indicate that Slx8 plays a critical role in regulating SUMO homeostasis in the nuclear space to safeguard centromere biology.

In several organisms, STUbL activities are linked to the maintenance of genome stability and resistance to DNA damage (reviwed in (Chang et al., 2021). In *S. pombe*, Slx8 operates with Ufd1, a component of the Cdc48-Udf1-Npl4 that allows the extraction of ubiquitylated proteins from higher-order complexes (Køhler et al., 2013). Both physical and functional overlaps between Ufd1 and Slx8 have revealed that Slx8 helps in channeling SUMOylated proteins towards such extraction process. This mechanism is part of the DNA damage response as Ufd1 forms DNA damage-induced foci, co-localizing with SUMO at the nuclear periphery. Similarly, ScSlx5 forms damage-induced nuclear foci in a SUMO-dependent manner, and co-localizing with DNA repair factors (Cook et al., 2009). Therefore, it was unanticipated that DNA damage does not lead to a redistribution of Slx8 to form specific DNA repair-associated foci. One possibility is that the amount of Slx8 recruited at site of DNA damage is below the level of detection offered in our cell microscopy condition. Alternatively, DNA-damage induced Slx8 foci are dynamic and rapidly moving to the SPB, leading to the increased foci intensity that we observed without increasing foci number.

Our observation of an Slx8 focus colocalizing with centromeres is consistent with several previous studies indicating that STUbLs reside and function at centromeres. In budding yeast, genome-wide binding analyses revealed centromeric enrichment of Slx5, but not of Slx8, and cells lacking Slx5 or Slx8 display chromosome segregation defects (van de Pasch et al., 2013). Indeed, Slx5-Slx8 STUbL activity has been shown to be required for degradation of several centromere-associated substrates including cohesion subunit Mcd1 (D’Ambrosio and Lavoie, 2014), chromosome passenger complex (CPC) components Bir1 and Sli15 (Thu et al., 2016), and centromere-specific histone H3 variant Cse4^CENP-A^ (Cheng et al., 2017; Ohkuni et al., 2018), thereby promoting the proper specification and function of centromeres. Similarly, mammalian RNF4 has been implicated in regulating centromere and kinetochore assembly, functioning antagonistically with SUMO protease SENP6 to modulate levels of CENP-A assembly factor Mis18BP1 (Fu et al., 2019; Liebelt et al., 2019) and inner kinetochore protein CENP-I (Mukhopadhyay et al., 2010). In *S. pombe*, it has been reported previously that loss of Slx8 results in chromosome segregation defects, dependent on the SUMO ligase Pli1 (Steinacher et al., 2013); our findings that absence of Slx8 is associated with defects in both heterochromatic silencing and centromere clustering point to multifaceted roles of STUbL activity in supporting normal centromere function.

Heterochromatin is a key structural and regulatory component of centromeres in most eukaryotes, functioning to promote accurate chromosome segregation and silence repetitive DNA elements. Perturbation of SUMOylation has been linked to defects in heterochromatic silencing in several systems, including flies (Ninova et al., 2020), mammals (Marshall et al., 2010), and *S. pombe* (Shin et al., 2005). Moreover, large-scale studies in various organisms have identified heterochromatic regions including centromeres as SUMOylation hotspots (Cubeñas-Potts and Matunis, 2013; Ninova et al., 2023). Indeed, in *S. pombe*, proteomic analyses revealed that more than a third of SUMOylated proteins regulated by Slx8 and Ufd1 are proteins associated with centromeres or telomeres, including key heterochromatin regulators, the H3K9 methyltransferase Clr4 and anti-silencing factor Epe1 (Køhler et al., 2015). How SUMOylation impacts the function of these specific proteins is yet to be established. However, it has previously been shown that Epe1 is subject to ubiquitin-dependent cleavage and degradation that regulates its activity within heterochromatin domains (Braun et al., 2011) and in response to stress (Yaseen et al., 2022); it is tempting to speculate that Slx8 STUbL activity might contribute to Epe1 ubiquitination, and therefore that alleviation of heterochromatic silencing in *slx8Δ* cells could potentially be attributable, at least in part, to increased Epe1 activity. Since the localization of potential substrates such as Epe1 and Clr4 at centromeres and the *mat* locus is heterochromatin-dependent (Isaac et al., 2007; Zofall and Grewal, 2006), this could help explain why Slx8 association with these regions is both dependent on, and required for, proper heterochromatin maintenance.

The phenomenon of centromere clustering has been observed in many eukaryotes and also appears to be important for normal centromere function, although the underlying mechanisms are not yet fully understood. In fission yeast, Csi1 plays an important role in tethering kinetochores to the SPB, and loss of this protein results in centromere declustering (Hou et al., 2012). We previously uncovered a role for SUMOylation in enhancing centromere clustering in conditions where Csi1 is absent, since removal of nucleoporin Nup132, which tethers SUMO protease Ulp1 to the NP, causes a SUMO-dependent rescue of clustering in *csi1Δ* cells (Strachan et al., 2023). This effect was found to be dependent on SUMOylation of the inner nuclear membrane protein Lem2, which acts in parallel with Csi1 to promote clustering, but independent of Slx8 activity. In contrast, here we show that Slx8 is required to maintain proper centromere clustering in otherwise wild-type cells. The relevant substrate(s) in this case are yet to be determined; however, our genetic data suggests that substrate(s) likely lie in the same pathway as Csi1, and therefore could potentially include, for example, Csi1 itself, or the interacting NE protein Sad1, both of which have been shown to be subject to SUMOylation (Køhler et al., 2015). In principle, Slx8 activity may be required either to temper the accumulation of SUMOylated proteins, or to actively promote protein extraction/turnover in a SUMO-dependent manner. However, we have shown previously that loss of the SUMO ligase Pli1 has only minimal effect on centromere clustering in this background (∼2.5% of *pli1Δ* cells displaying declustering, as compared to ∼12% of *slx8Δ* cells), whereas we confirm here that the clustering defects associated with absence of Slx8 are largely suppressed upon removal of Pli1. Thus, it is likely that Slx8 is primarily required to prevent the detrimental excess accumulation of SUMOylated substrates, and therefore to help maintain an optimal balance of SUMOylation needed to support normal centromere clustering.

SUMOylation has been found to influence the dynamics of telomere maintenance in *S. pombe* by controlling the activity of positive or negative regulators of telomerase (Xhemalce et al., 2004; Xhemalce et al., 2007). In our microscopy analysis, we could not detect association between Slx8 and Taz1-marked telomeres. It is worth noting that, in budding yeast, telomeric factors are enriched for SUMO modifications upon telomere erosion (in the absence of telomerase), resulting in Slx5/Slx8-mediated relocation to the NPC to promote telomere length maintenance (Churikov et al., 2016). Therefore, it is possible that, without stress-inducing SUMOylation at telomeres, the association between Slx8 and telomere is below limit of detection. Nonetheless, our results highlight that, in unchallenged conditions, Slx8 mainly acts at heterochromatic domains and centromeres to orchestrate the nuclear organization and functions of these specific domains.

## Materials and methods

### Standard yeast genetics and biological resources

Yeast strains used in this work are listed in Table S1. Gene deletion and tagging were performed by classical genetic techniques. To assess the sensitivity of chosen mutants to genotoxic agents, mid log-phase cells were serially diluted and spotted onto yeast extract agar plates containing hydroxyurea (HU), methyl methanesulfonate (MMS), campthotecin (CPT).

### Live cell imaging

For snapshot microscopy, cells were grown in filtered EMMg to exponential phase, then centrifuged and resuspended in 500 µL of fresh EMMg. 1 µL from the resulting solution was dropped onto Thermo Scientific slide (ER-201B-CE24) covered with a thin layer of 1.4 % agarose in filtered EMMg (Kramarz et al., 2020). 11 z-stack pictures (each z step of 200 nm) were captured using a Nipkow Spinning Disk confocal system (Yokogawa CSU-X1-A1) mounted on a Nikon Eclipse Ti E inverted microscope, equipped with a 100x Apochromat TIRF oil-immersion objective (NA: 1.49) and captured on sCMOS Prime 95B camera (Photometrics) operated through MetaMorph^®^ software (Molecular Devices). The GFP proteins were excited with a 488 nm (Stradus® - Vortran Laser Technology, 150mW) laser, while RFP and m-Cherry proteins were excited with a 561 nm (Jive^TM^ - Cobolt, 100mW) laser. A quad band dichroic mirror (405/488/568/647 nm, Semrock) was used in combination with single band-pass filters of 525/50 or 630/75 for the detection of GFP, RFP and m-Cherry, respectively. Fluorescence and bright-field 3D images were taken at every 0.2µm by acquiring one wavelength at a time. Exposure time for GFP channel was 500 ms, for RFP was 300 ms and for mCherry was 600 ms. During the imaging, the microscope was set up at 25°C. For all the experiments the Gataca Live SR module (Müller et al., 2016, Gataca Systems), implemented on the Spinning Disk confocal system, was used to generate super-resolution images. All image acquisition was performed on the PICT-IBiSA Orsay Imaging facility of Institut Curie.

### Image analysis

Images were mounted and analyzed with Fiji software (Schindelin et al., 2012). First, the 3D Z series were converted into 2D projection based on maximum intensity values to produce the image with merged stacks. Since, Slx8 is a low abundant protein, with a high nuclear background, the quantification of Slx8 foci were performed using a noise tolerance threshold value of 50 (Maxima) from Fiji. This was decided after comparing different Maxima values in order to detect foci vs random background noise. Once the threshold was applied, the foci could be manually counted by selecting them as detected by the software. All experiments have been analysed with the same Maxima value in this report. For quantification of the percentage of co-localization between Slx8 and other markers, the same as above was done onto the GFP channel to first annotate the Slx8 foci above the “set” threshold. In a separate window, the GFP and RFP/mCherry channels with different stacks were merged together followed by manually analysing the co-localization of the green and red foci signal at each stack. Maxima was not applied to RFP/mCherry channel because the foci detection was clear and obvious with no nuclear background noise. The probability of a random event for the co-localisation experiments were performed by using the 180° transform tool in Fiji for the RFP/mCherry marker, followed by merge with the normal Slx8 GFP stacks (without the 180° transform). Consequently, analysis of co-localization between the green and red foci signal at each stack in this setting provided the number of random co-localisation events possible in each given field. This value is referred to as the “random event” that provides a threshold to calculate the possibility of significant co-localisation events as compared to random events.

### Centromere clustering analysis

For clustering analysis, cells expressing GFP–Cnp1 and Sid4–RFP were grown in YES to exponential phase, then centrifuged and resuspended in 30 µL YES. 4 uL of the resulting cell suspension was mixed with 6 uL of 1% low-melting point agarose and imaging was performed at 25°C using a Nikon Ti2 inverted microscope, equipped with a 100×1.49 NA Apo TIRF objective and a Teledyne Photometrics Prime 95B camera. Images were acquired with NIS-elements (version 5.1), with *Z*-stacks taken at 250 nm intervals. Maximum intensity *Z*-projections were made in ImageJ. Manual quantification of the number of GFP foci per cell was performed to determine the proportions of cells displaying centromeres ‘clustered’ (one GFP–Cnp1 focus) versus ‘unclustered’ (two or three GFP–Cnp1 foci).

### Whole protein extract analysis

Aliquots of 1×10^8^ cells were collected and disrupted by bead beating in 1 mL of 20 % TCA (Sigma, T9159). Pellets of denatured proteins were washed with 1M Tris pH 8 and resuspended in 2x Laemmli buffer (62.5 mM Tris pH 6.8, 20 % glycerol, 2 % SDS, 5 % β-mercaptoethanol with bromophenol blue). Samples were boiled before being subjected to SDS-PAGE on Mini-PROTEAN TGX Precast Gel 4-15 % (Biorad, 4561086). Western blot using anti-GFP (Roche, 11814460001) and anti-PCNA (Santa Cruz, sc-56) antibodies was performed. For the analysis of cellular patterns of global SUMOylation, whole protein extraction was performed as follows: aliquots of 2×10^8^ cells were collected and resuspended in 400µl of water. The cell suspensions were mixed with 350 µl of freshly prepared lysis buffer (2M NaOH, 7% β-mercaptoethanol) and 350µl of 50% TCA (Sigma, T9159). After spin, pellets were further washed with 1M Tris pH 8 and resuspended in 2x Laemmli buffer (62.5 mM Tris pH 6.8, 20 % glycerol, 2 % SDS, 5 % β-mercaptoethanol with bromophenol blue). Samples were boiled before being subjected to SDS-PAGE on Mini-PROTEAN TGX Precast Gel 4-15 % (Biorad, 4561086). Western blot using anti-SUMO antibody (non-commercial, produced in rabbit by Agro-Bio) was performed (dilution of 1:1000).

### RT-qPCR

Total RNA was extracted from 1×10^7^ mid-log phase cells using the Masterpure Yeast RNA Purification Kit (Epicentre), according to the manufacturer’s instructions. 1 µg of extracted RNA was treated with TURBO DNase (Ambion) for 1 h at 37°C, and reverse transcription was performed using random hexamers (Roche) and Superscript III reverse transcriptase (Invitrogen). Lightcycler 480 SYBR Green (Roche) and primers (_q_cen[dg]_F: 5’-AATTGTGGTGGTGTGGTAATAC-3’ and _q_cen[dg]_R: 5’-GGGTTCATCGTTTCCATTCAG-3’; _q_mat[D]_F: 5’-GTCCGAGGCAATACAACTTTGG-3’; and _q_mat[D]_R: 5’-GGTTGACAGTAGGAGATATTTACAG-3’; _q_act1_F: 5′-GTTTCGCTGGAGATGATG-3′ and _q_act1_R: 5′-ATACCACGCTTGCTTTGAG-3′) were used for qPCR quantification of pericentromere (*dg*) and mating type locus (*mat*) transcript levels, relative to *act1^+^*.

## STATISTICAL ANALYSIS

Quantitative analysis of western blots were carried out using Fiji software. The ratio from the Raw Integrated Density value of the protein of interest to housekeeping control was calculated for estimating the amount of protein.

Cell imaging was performed using METAMORPH software and processed and analyzed using ImageJ software (ref). The explanation and definitions of values and error bars are mentioned within the figure legends. In most experiments, the number of samples is > 2 and obtained from independent experiments to ensure biological reproducibility. For all experiments based on the analysis of cell imaging, the number of nuclei analyzed is mentioned in the figure legends. Statistical analysis was carried out using Mann-Whitney U tests, Brown-Forsythe and Welch Anova test, chi-squared test and Student’s *t*-test. ns: P≥ 0.05, *P≤ 0.05, **P ≤ 0.01, ***P≤ 0.001, ****P ≤ 0.0001.

## Acknowledgements

The authors thank the Multimodal Imaging Center Imaging Facility of the Institut Curie - CNRS UMS2016 / Inserm US43 / Institut Curie / Université Paris-, and the Centre Optical Instrumentation Laboratory (COIL) at the University of Edinburgh, supported by the Wellcome Trust (203149). We are also grateful to Robin Allshire, Yasushi Hiraoka and Matthew Whitby for sharing yeast strains.

S.C., J.S., K.S., E.V., K.F, H.Z. and N.Z. performed the experiments.

S.C., J.S., K.S., E.H.B. and S.A.E.L contributed to experimental design and data analysis.

L.B. provided expertise to perform and analyze cell imaging.

E.H.B. and S.A.E.L wrote the manuscript.

## Data and resource availability

Source data files containing raw data are deposited at Mendeley data under doi: 10.17632/tf585hwkj3.1.

All relevant data are available and further information and requests for reagents and resources should be directed to and will be fulfilled by Dr. Sarah A.E. Lambert.

## Supplemental Information

**Figure S1:**
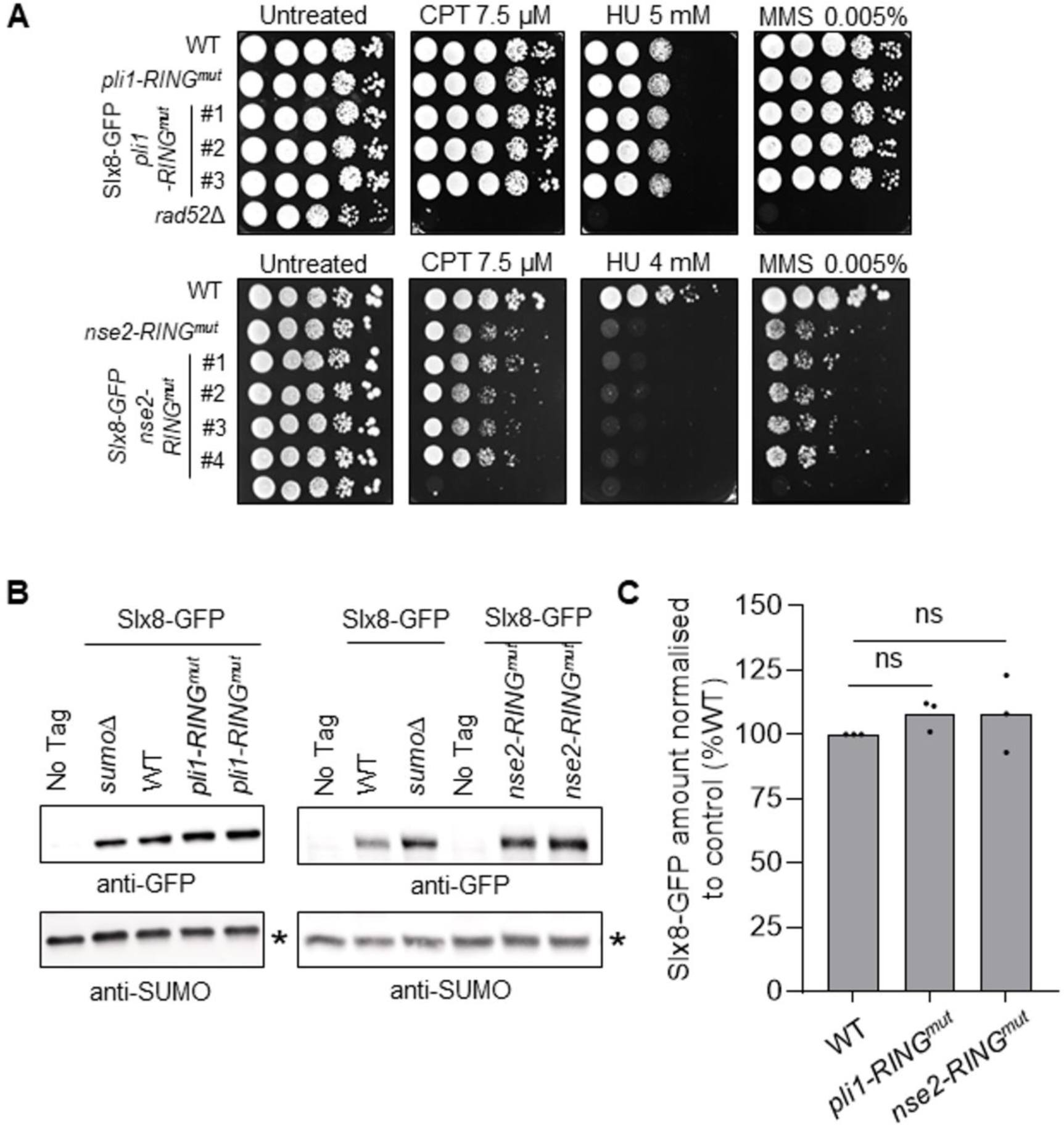
Expression of Slx8-GFP is not affected by the absence of the E3 SUMO ligase activity of either Nse2 or Pli1. **A.** Sensitivity of indicated strains to genotoxic drugs. Ten-fold serial dilution of exponential cultures were dropped onto indicated plates. HU: hydroxyurea; CPT: camptothecin and MMS: methyl methane sulfonate. **B.** Expression of Slx8-GFP in indicated strains. An untagged WT strain (No Tag) was included as control for antibody specificity. An unspecific band (*) from SUMO-blots was used as a loading control. **C.** Quantification of Slx8-GFP expression in indicated strains. Dots represent values obtained from independent biological experiments. The normalized amount of Slx8 was calculated by dividing the GFP signal by unspecific SUMO signal. The normalized amount of Slx8-GFP in mutants was indicated as a percentage of WT. *p* value was calculated by two-sided Fisher’s exact test (ns: non-significant).

**Figure S2:**
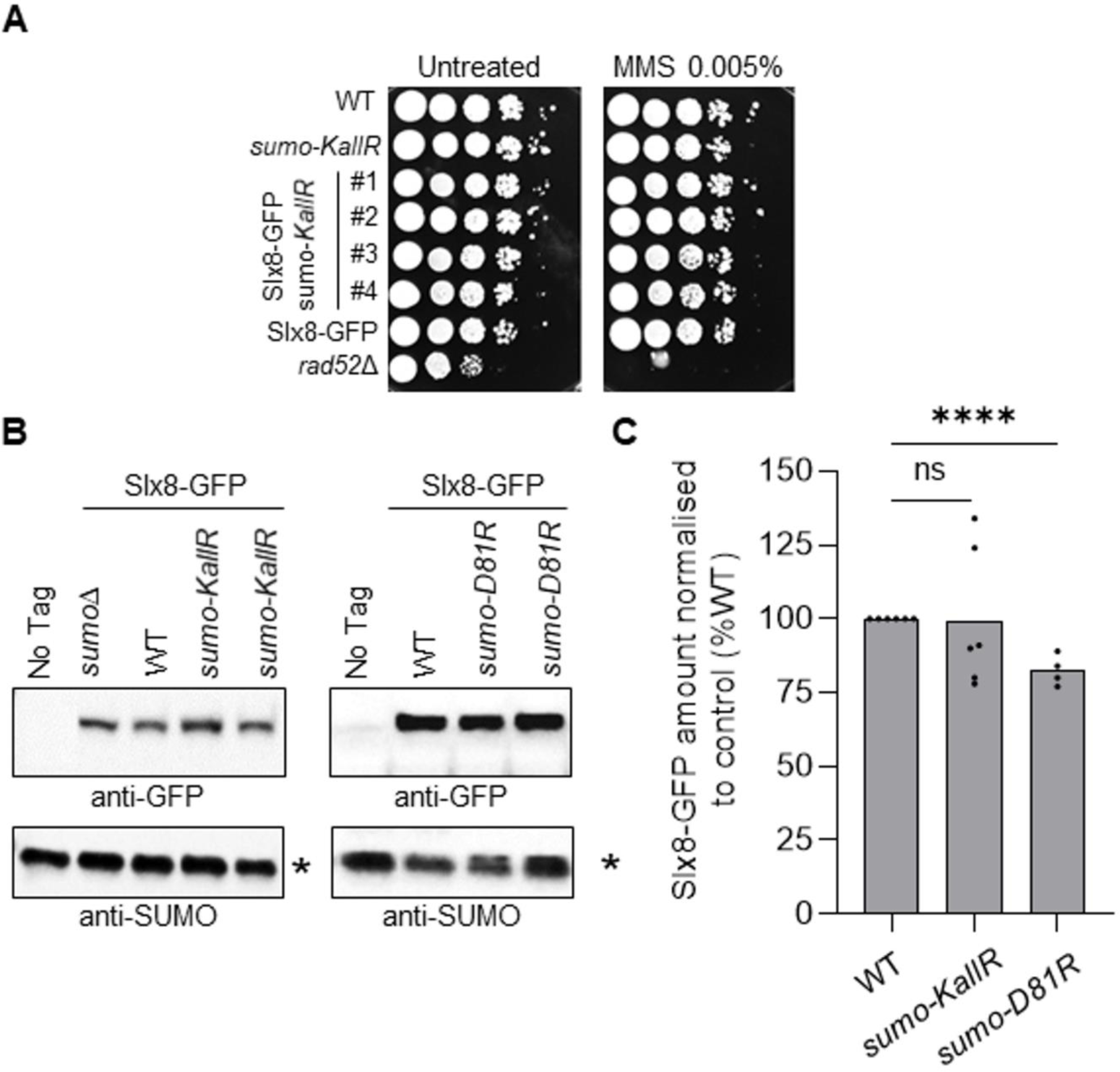
Expression of Slx8-GFP is not affected in strains expressing SUMO-D81R or SUMO-KallR. **A.** Sensitivity of indicated strains to genotoxic drugs. Ten-fold serial dilution of exponential cultures were dropped onto indicated plates. MMS: methyl methane sulfonate. **B.** Expression of Slx8-GFP in indicated strains. An untagged WT strain (No Tag) was included as control for antibody specificity. An unspecific band (*) from SUMO-blots was used as a loading control. **C.** Quantification of Slx8-GFP expression in indicated strains. Dots represent values obtained from independent biological experiments. The normalized amount of Slx8 was calculated by dividing the GFP signal by unspecific SUMO signal. The normalized amount of Slx8-GFP in mutants was indicated as a percentage of WT. *p* value was calculated by two-sided Fisher’s exact test (**** p<0.0001; ns: non-significant).

**Figure S3:**
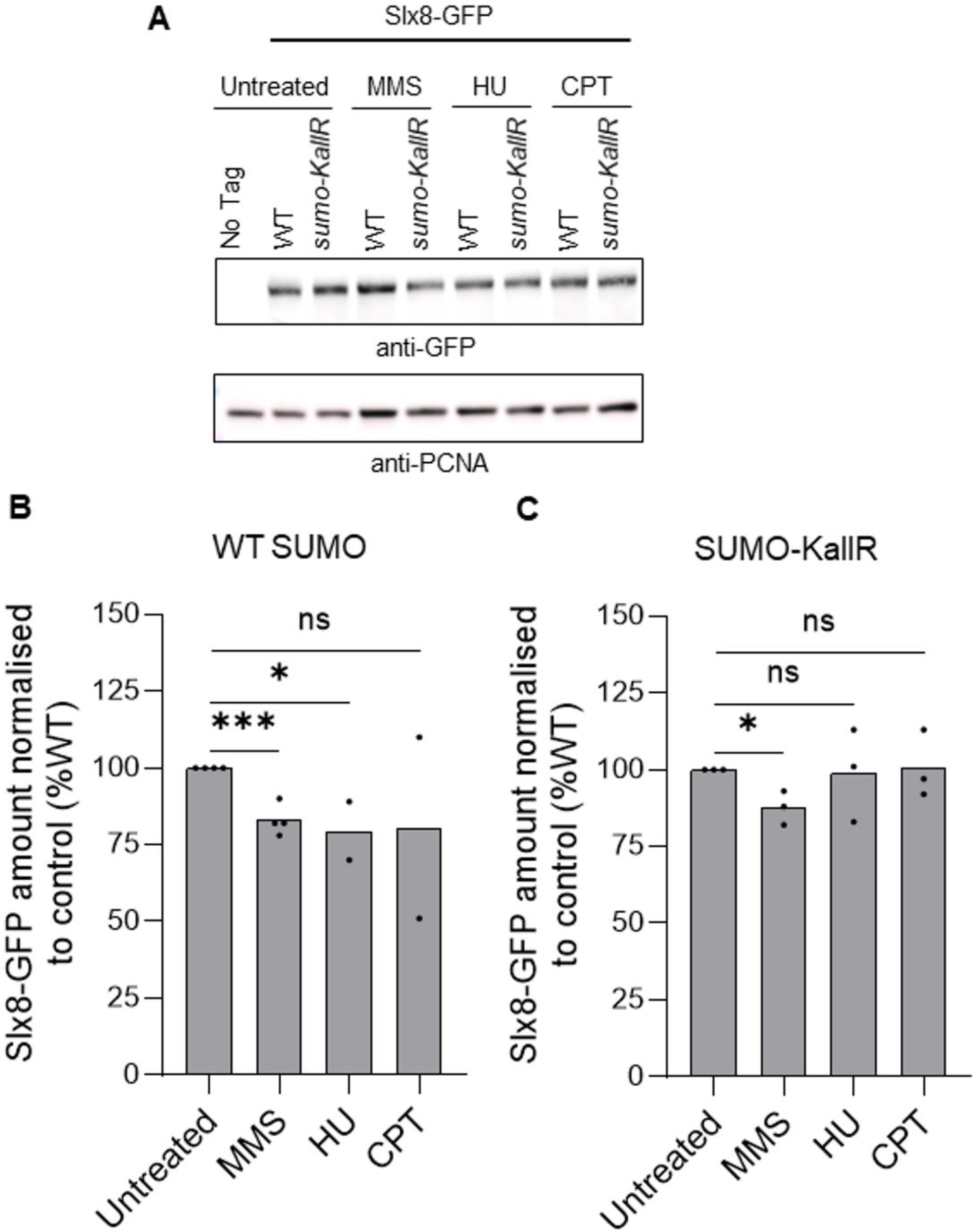
Genotoxic treatments have a variable effect upon Slx8-GFP protein expression profile. **A.** Expression of Slx8-GFP in indicated strains and conditions. An untagged WT strain (No Tag) was included as control for antibody specificity. PCNA was used as a loading control. HU: hydroxyurea; CPT: camptothecin and MMS: methyl methane sulfonate. **B & C.** Quantification of Slx8-GFP expression in indicated strains (WT SUMO: left panel, SUMO-KallR: right panel) and conditions. Dots represent values obtained from independent biological experiments. The normalized amount of Slx8 was calculated by dividing the GFP signal by PCNA signal. The normalized amount of Slx8-GFP in treated conditions was indicated as a percentage of the untreated conditions. *p* value was calculated by two-sided Fisher’s exact test (*** p≤0.001; * p≤0.05; ns: non-significant).

**Figure S4:**
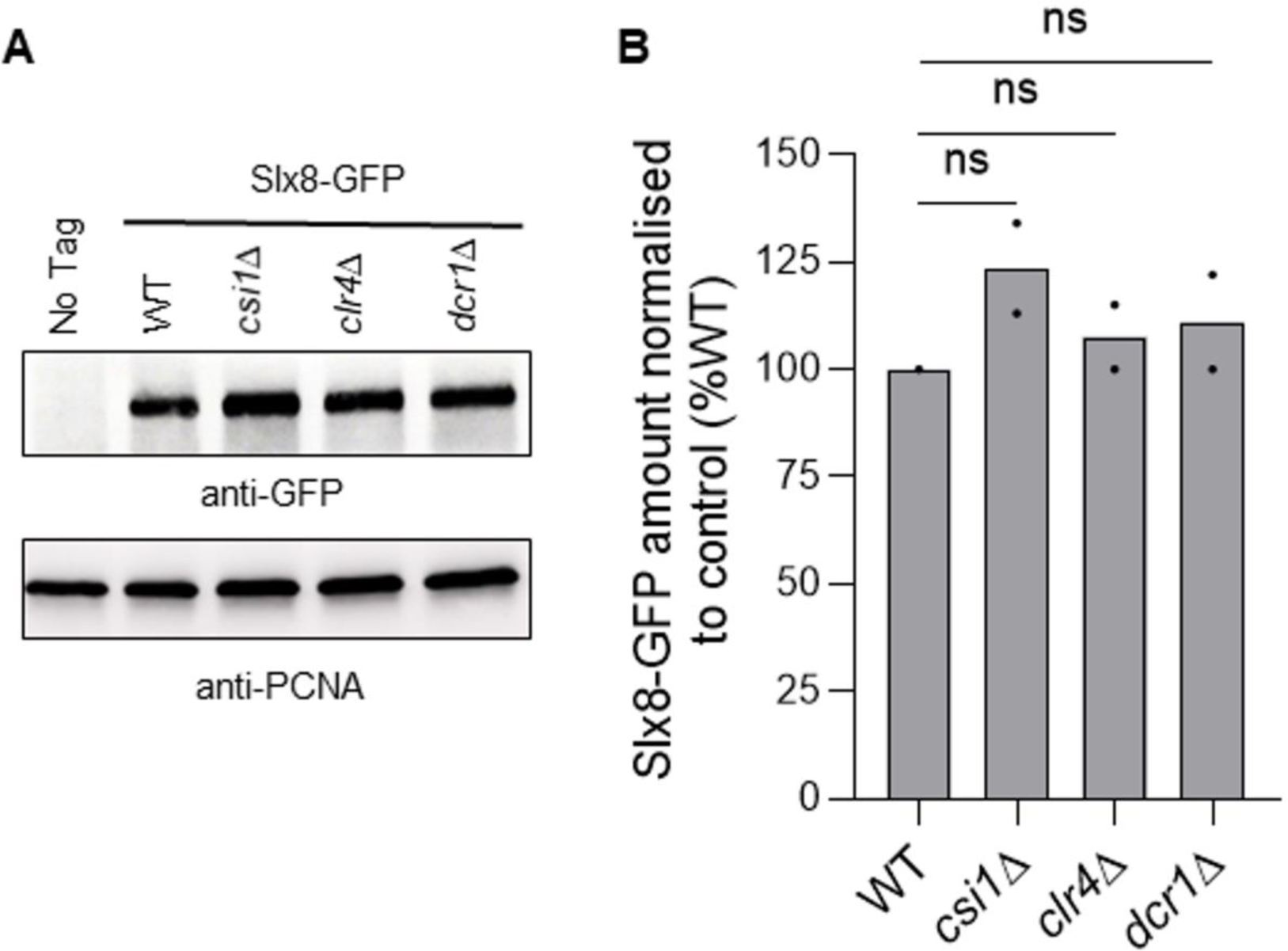
Expression of Slx8-GFP is not affected by the absence of Clr4, Dcr1 or Csi1. **A.** Expression of Slx8-GFP in indicated strains. An untagged WT strain (No Tag) was included as control for antibody specificity. PCNA was used as a loading control. **B.** Quantification of Slx8-GFP expression. Dots represent values obtained from independent biological experiments. The normalized amount of Slx8 was calculated by dividing the GFP signal by PCNA signal. The normalized amount of Slx8-GFP in mutants was indicated as a percentage of the WT. *p* value was calculated by two-sided Fisher’s exact test (ns: non-significant).

**Table S1:** Strains used in this study.

| Strain number | Mating type | Genotype | Reference |
| --- | --- | --- | --- |
| KK1492 | *h*- | *slx8-GFP:natMX6 nmt41:rtf1:sup35 ade6-704 leu1-32 t-ura4-SD20<ori (uraR)* | this study |
| KK2021 | *h*+ | *slx8-GFP:natMX6 ade6-704 leu1-32 ura4-D18* | this study |
| KK1377 | *h*+ | *ade6-704 leu1-32 ura4-D18* | this study |
| KK772 | *h*+ | *rad52::kanMX6 nmt41:rtf1:sup35 ade6-704 t-ura4<sup>+</sup><ori (uraR) leu1-32* | this study |
| KK1562 | *h*- | *pmt3::kanMX6-ura4<sup>+</sup> nmt41:rtf1:sup35 ade6-704 leu1-32 t-ura4-SD20<ori (uraR)* | this study |
| KK1025 | *h*- | *slx8-29:hphMX6 nmt41:rtf1:sup35 ade6-704 leu1-32 t-ura4-SD20<ori (uraR)* | this study |
| KK2096 | *h*- | *pli1-C321S-H323A-C326S (pli-RING<sup>mut</sup>) slx8-GFP:natMX6 ade6-704 leu1-32 ura4-D18* | this study |
| KK2112 | *h*+ | *nse2-C195S-H197A (nse2-RING<sup>mut</sup>) slx8-GFP:natMX6 ade6-704 leu1-32 ura4-D18* | this study |
| KK2023 | *h*+ | *pmt3-KallR (sumo-KallR) slx8-GFP:natMX6 ade6-704 leu1-32 ura4-D18* | this study |
| KK2074 | *h*- | *pmt3-D81R (sumo-D81R) slx8-GFP:natMX6 ade6-704 leu1-32 ura4-D18* | this study |
| KK2176 | *h*+ | *slx8-GFP:natMX6 cut11-mCherry:hphMX6 ade6-704 leu1-32 ura4-D18* | this study |
| KK2173 | *h*+ | *pmt3-KallR (sumo-KallR) slx8-GFP:natMX6 cut11-mCherry:hphMX6 ade6-704 leu1-32 ura4-D18* | this study |
| KK2294 | *h*- | *slx8-GFP:natMX6 sid4-mRFP:kanMX6 ade6-704 leu1-32 ura4-D18* | this study |
| KK2201 | *h*+ | *slx8-GFP:natMX6 mis6-mRFP:hphMX6 ade6-704 leu1-32 ura4-D18* | this study |
| KK2217 | *h*+ | *slx8-GFP:natMX6 taz1-mRFP:hphMX6 ade6-704 leu1-32 ura4-D18* | this study |
| KK2602 | *h90* | *slx8-GFP:natMX6 arg3::mCherry-LacI his2::kanR-ura4<sup>+</sup>-lacOp ade6-704 leu1-32 ura4-D18* | this study |
| KK2471 | *h*- | *csi1::hphMX6 slx8-GFP:natMX6 ade6-704 leu1-32 ura4-D18* | this study |
| KK2432 | *h*+ | *clr4::natMX6 slx8-GFP:natMX6 ade6-704 leu1-32 ura4-D18* | this study |
| KK2436 | *h-* | *dcr1::hphMX6 slx8-GFP:natMX6 ade6-704 leu1-32 ura4-D18* | this study |
| 673 | *h90* | *mat3-M:ade6<sup>+</sup> ade6-DN/N leu1-32 ura4-D18* | this study |
| 674 | *h90* | *clr4Δ::leu2 mat3-M:ade6<sup>+</sup> ade6-DN/N leu1-32 ura4-D18* | this study |
| 6711 | *h90* | *slx8Δ::ura4<sup>+</sup> mat3-M:ade6<sup>+</sup> ade6-DN/N leu1-32 ura4-D18* | this study |
| 5513 | *h90* | *Sid4<sup>+</sup>-mRFP:Kan<sup>R</sup> GFP-Cnp1<sup>+</sup>:Nat<sup>R</sup> leu1-32 ura4-D18* | this study |
| 6363 | *h+* | *csi1Δ::Hyg<sup>R</sup> Sid4<sup>+</sup>-mRFP:Kan<sup>R</sup> GFP-Cnp1<sup>+</sup>:Nat<sup>R</sup> leu1-32 ura4-D18* | this study |
| 7681 |  | *slx8Δ::Kan<sup>R</sup> Sid4<sup>+</sup>-mRFP:Kan<sup>R</sup> GFP-Cnp1<sup>+</sup>:Nat<sup>R</sup> leu1-32 ura4-D18* | this study |
| 8000 |  | *csi1Δ::Hyg<sup>R</sup> slx8Δ::Kan<sup>R</sup> Sid4<sup>+</sup>-mRFP:Kan<sup>R</sup> GFP-Cnp1<sup>+</sup>:Nat<sup>R</sup> leu1-32 ura4-D18* | this study |
| 8028 |  | *slx8Δ::Kan<sup>R</sup> pli1Δ::ura4<sup>+</sup> Sid4<sup>+</sup>-mRFP:Kan<sup>R</sup> GFP-Cnp1<sup>+</sup>:Nat<sup>R</sup> leu1-32 ura4-D18* | this study |
| 7036 |  | *pli1Δ::ura4<sup>+</sup> Sid4<sup>+</sup>-mRFP:Kan<sup>R</sup> GFP-Cnp1<sup>+</sup>:Nat<sup>R</sup> leu1-32 ura4-D18* | this study |

